## Supplementary figures and images for "A progeria-associated BAF-1 mutation modulates gene expression and accelerates aging in *C. elegans*"

### Supplementary Figures

Supplementary Figure 1

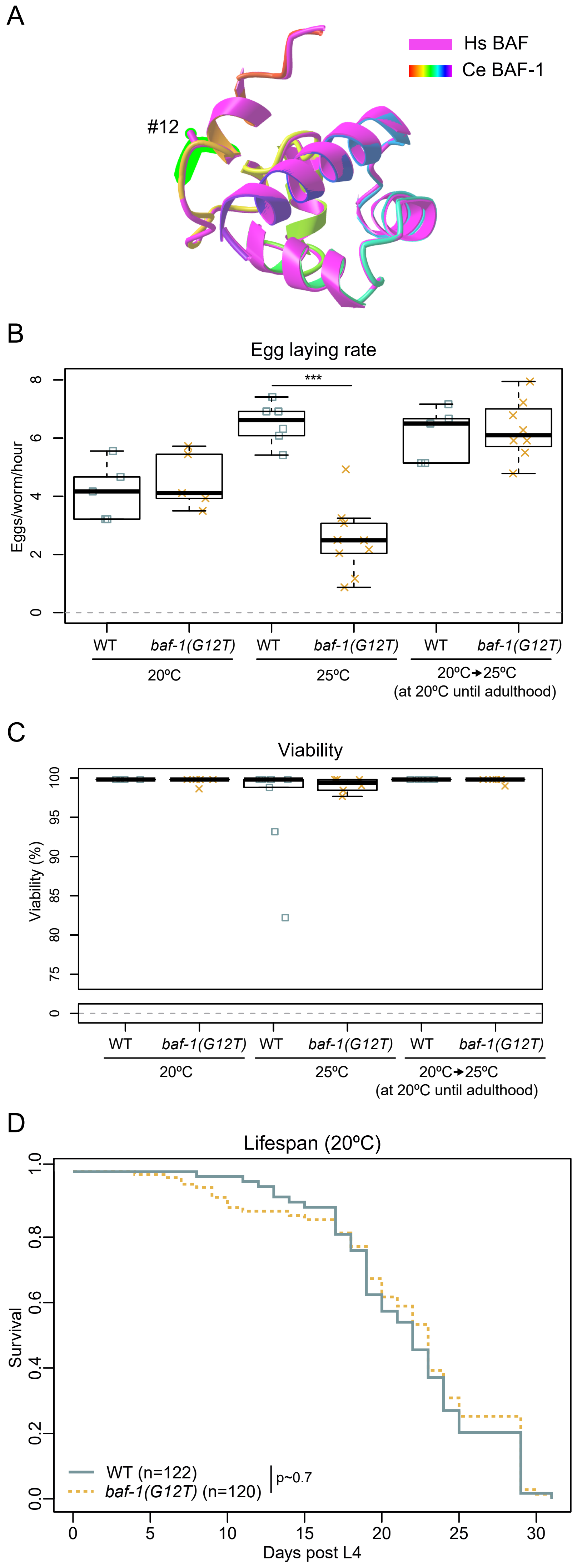

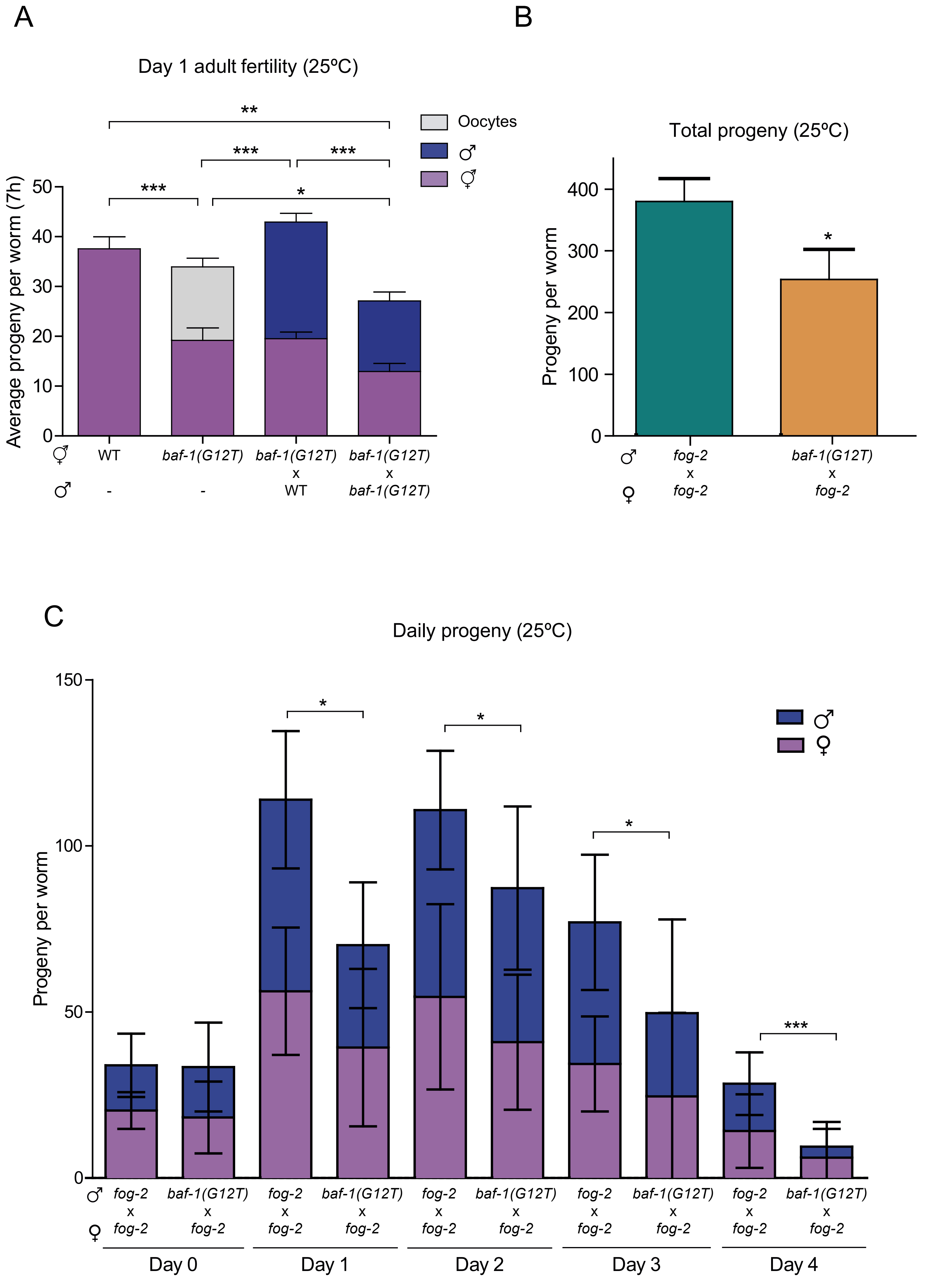

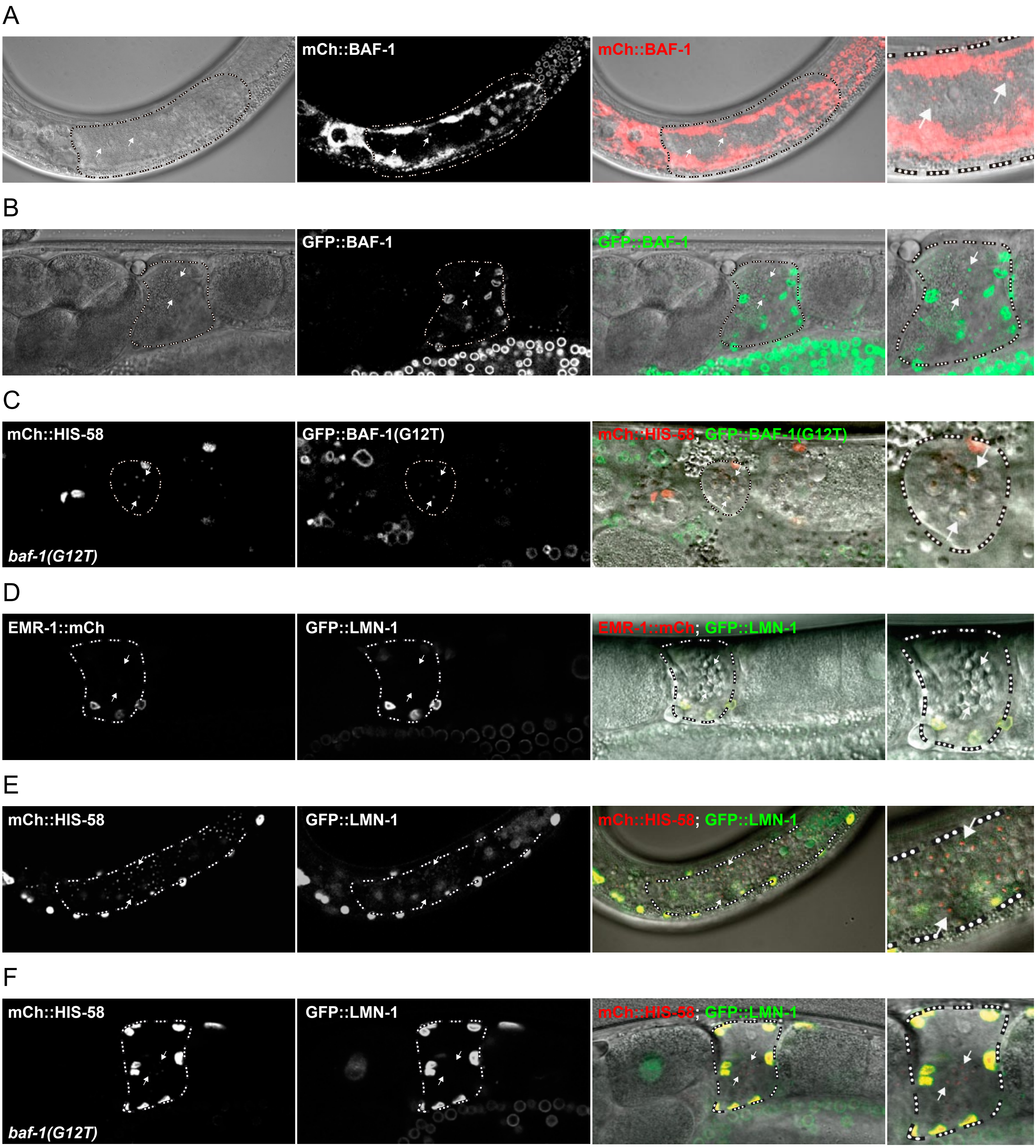

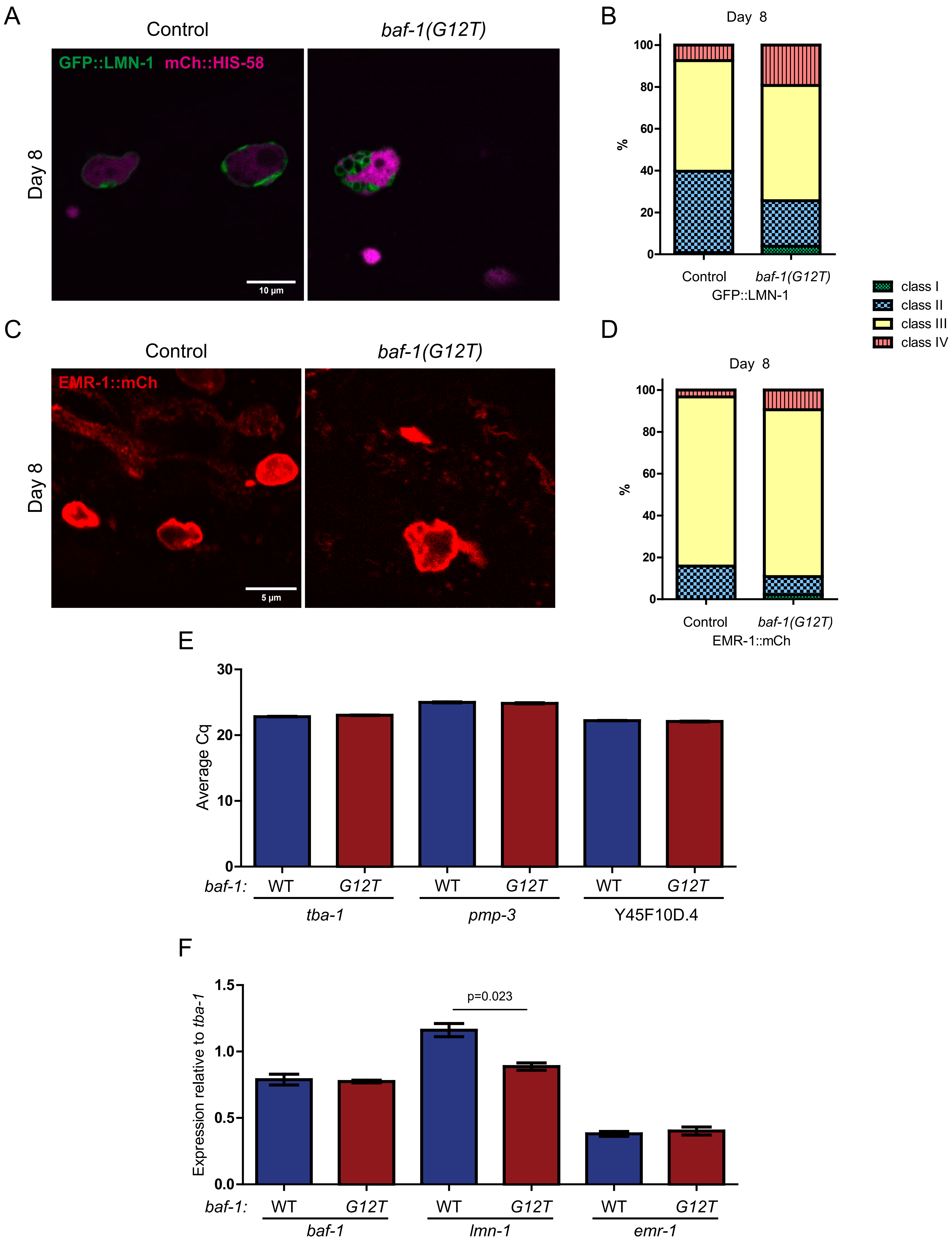

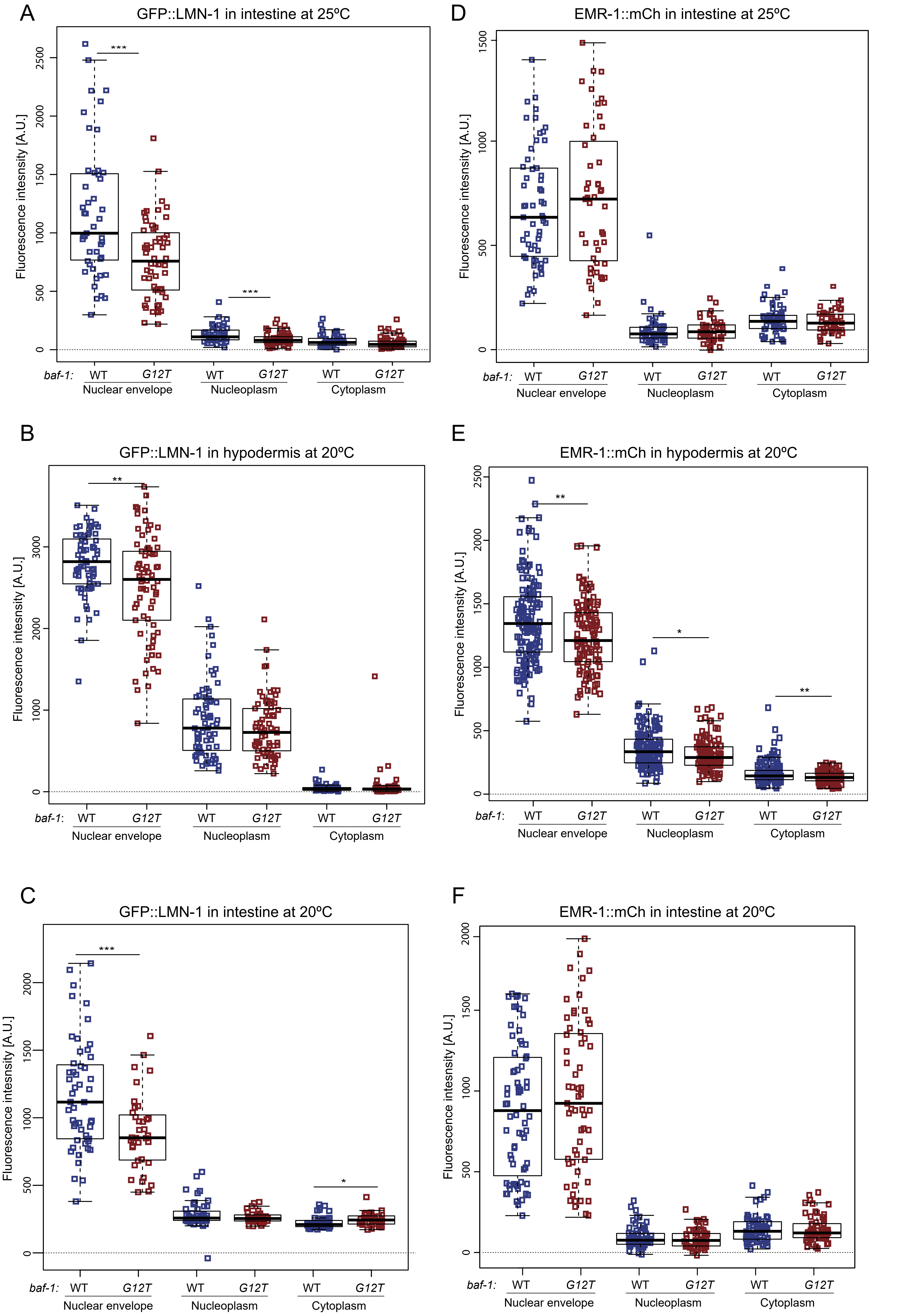

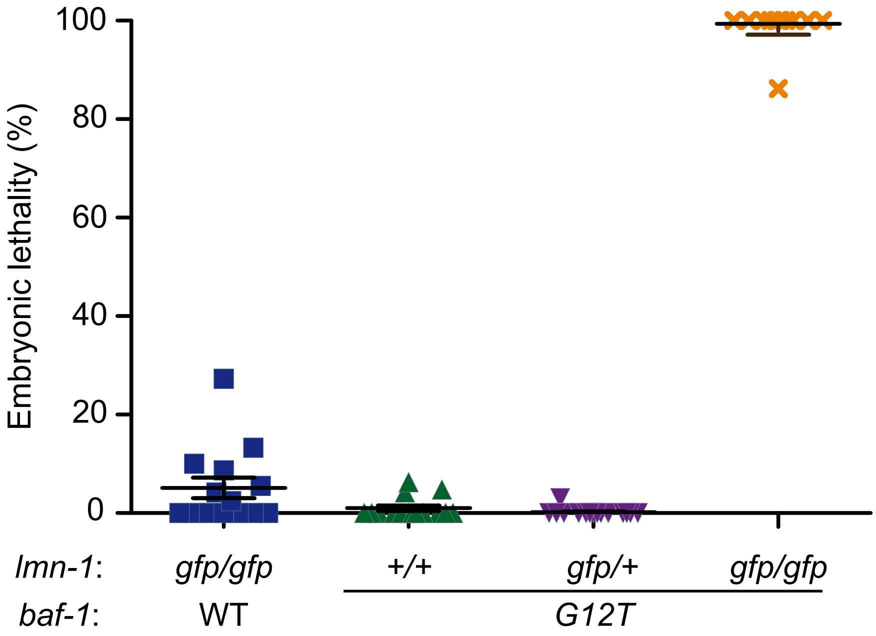

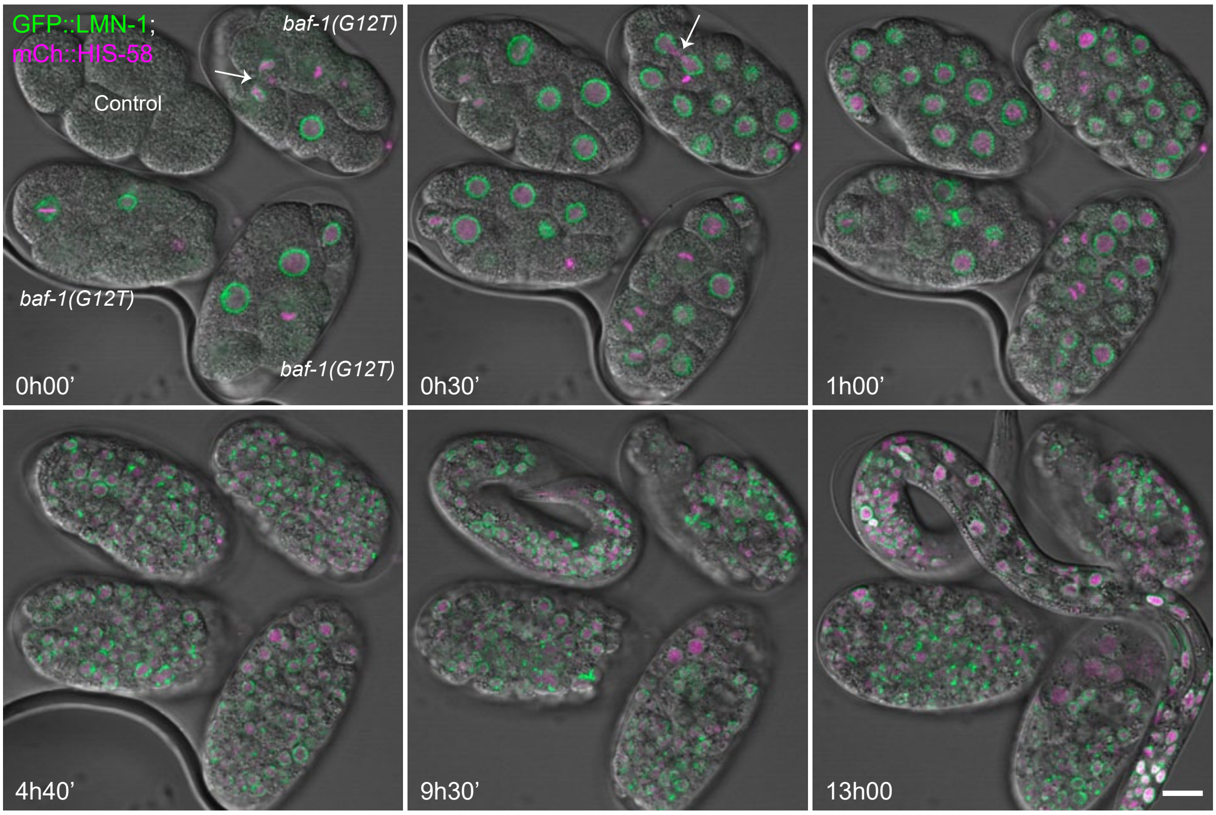

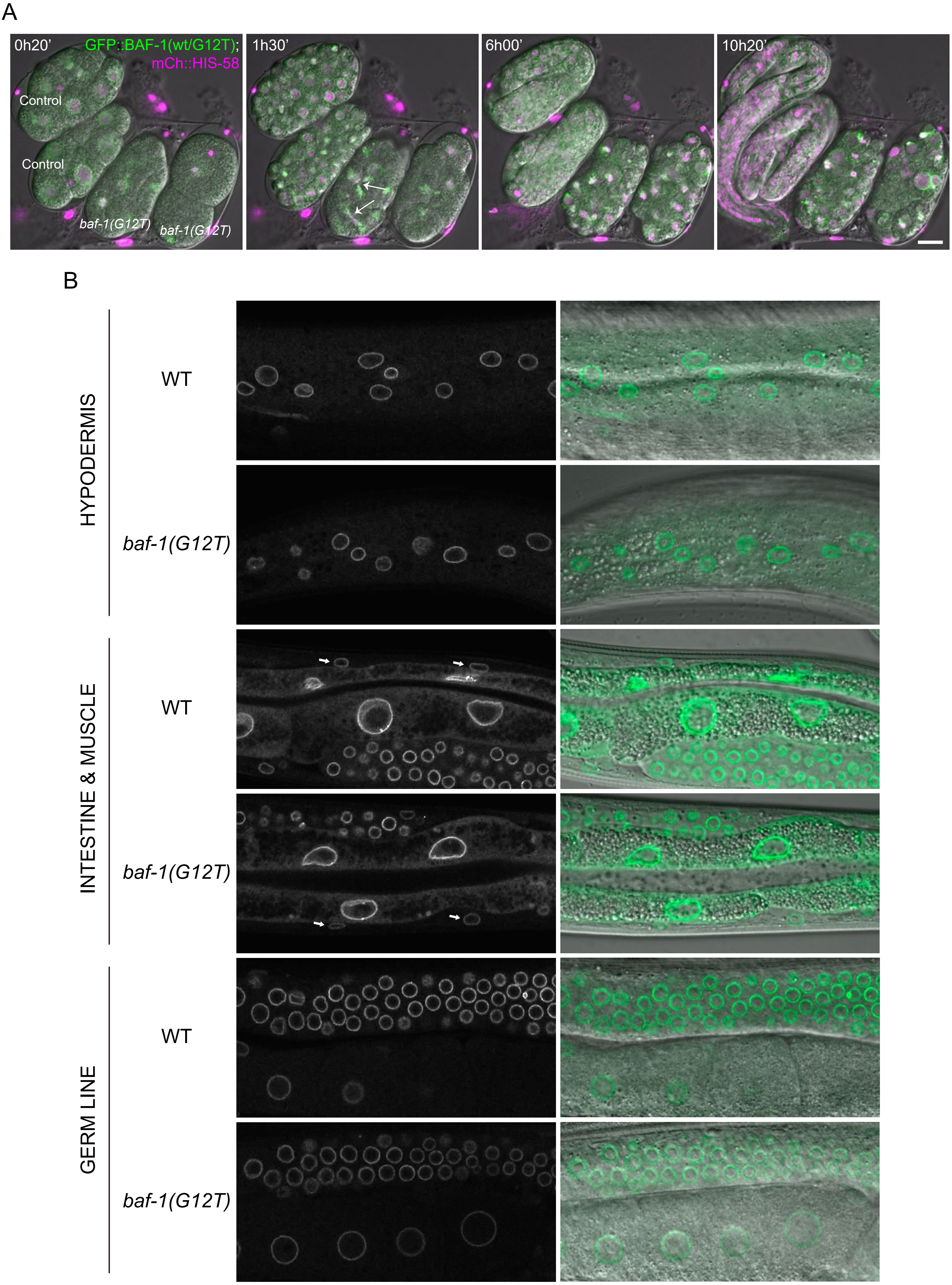

Supplementary Figure 8

A

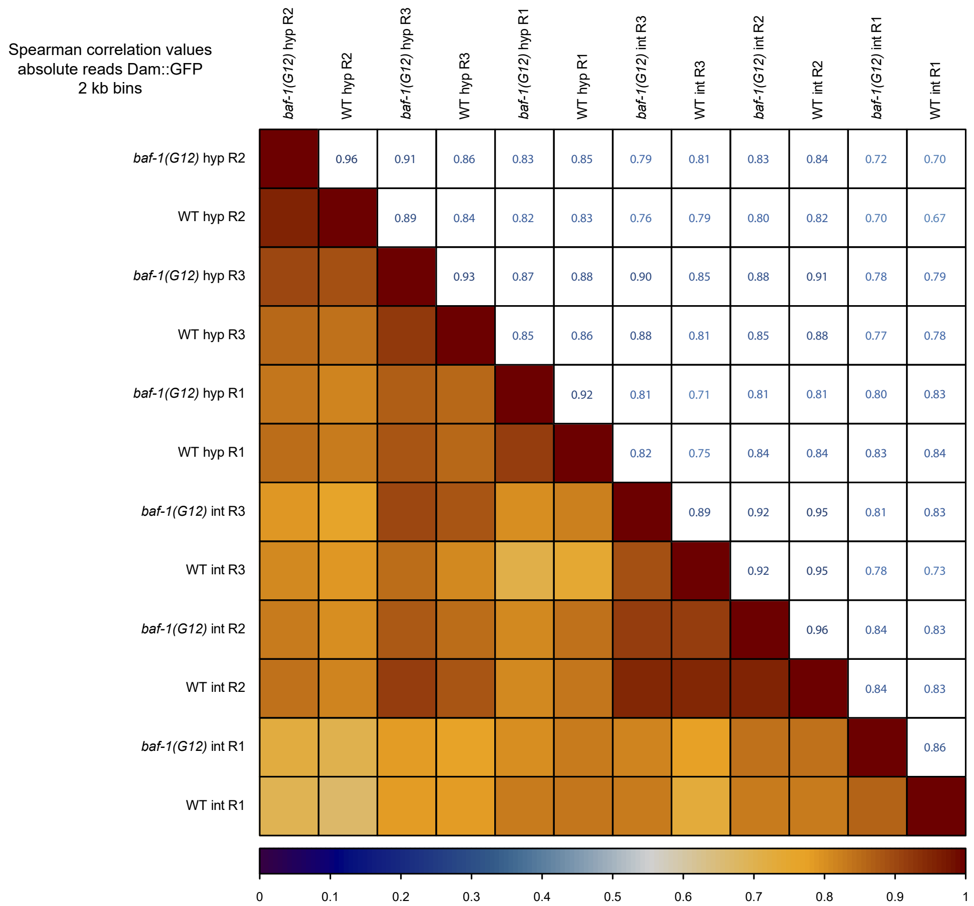


B

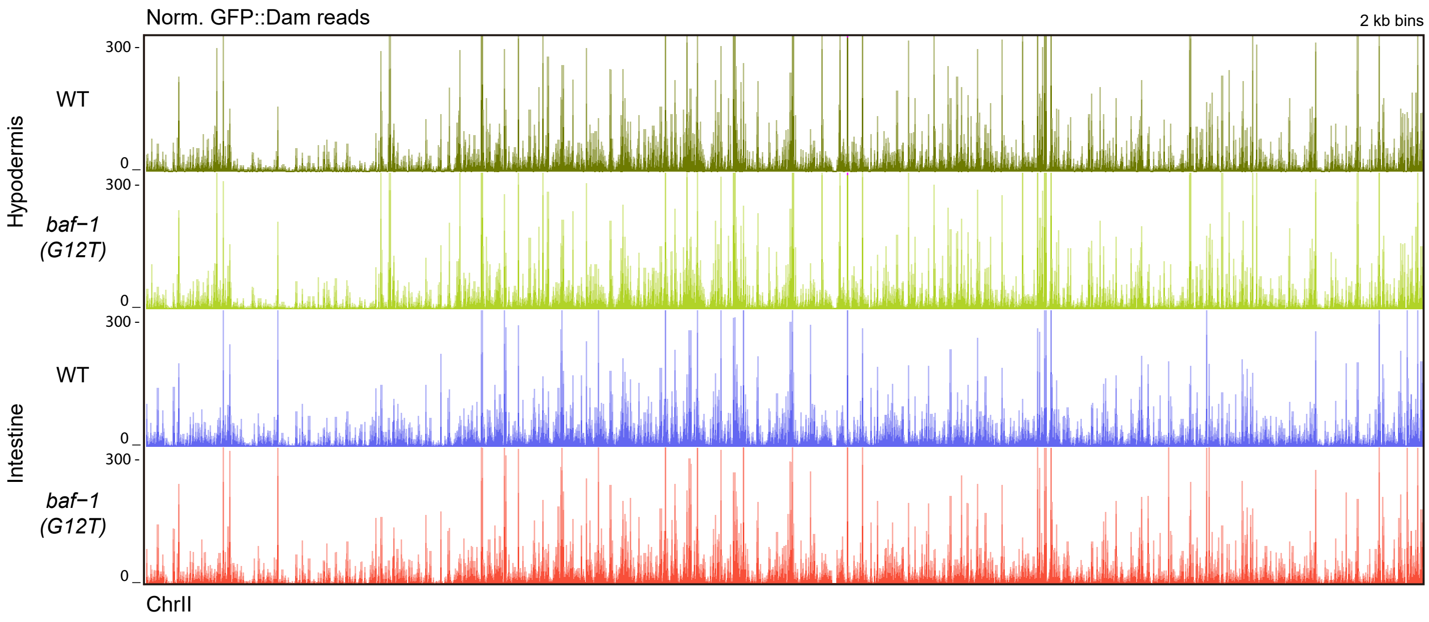

Supplementary Figure 9

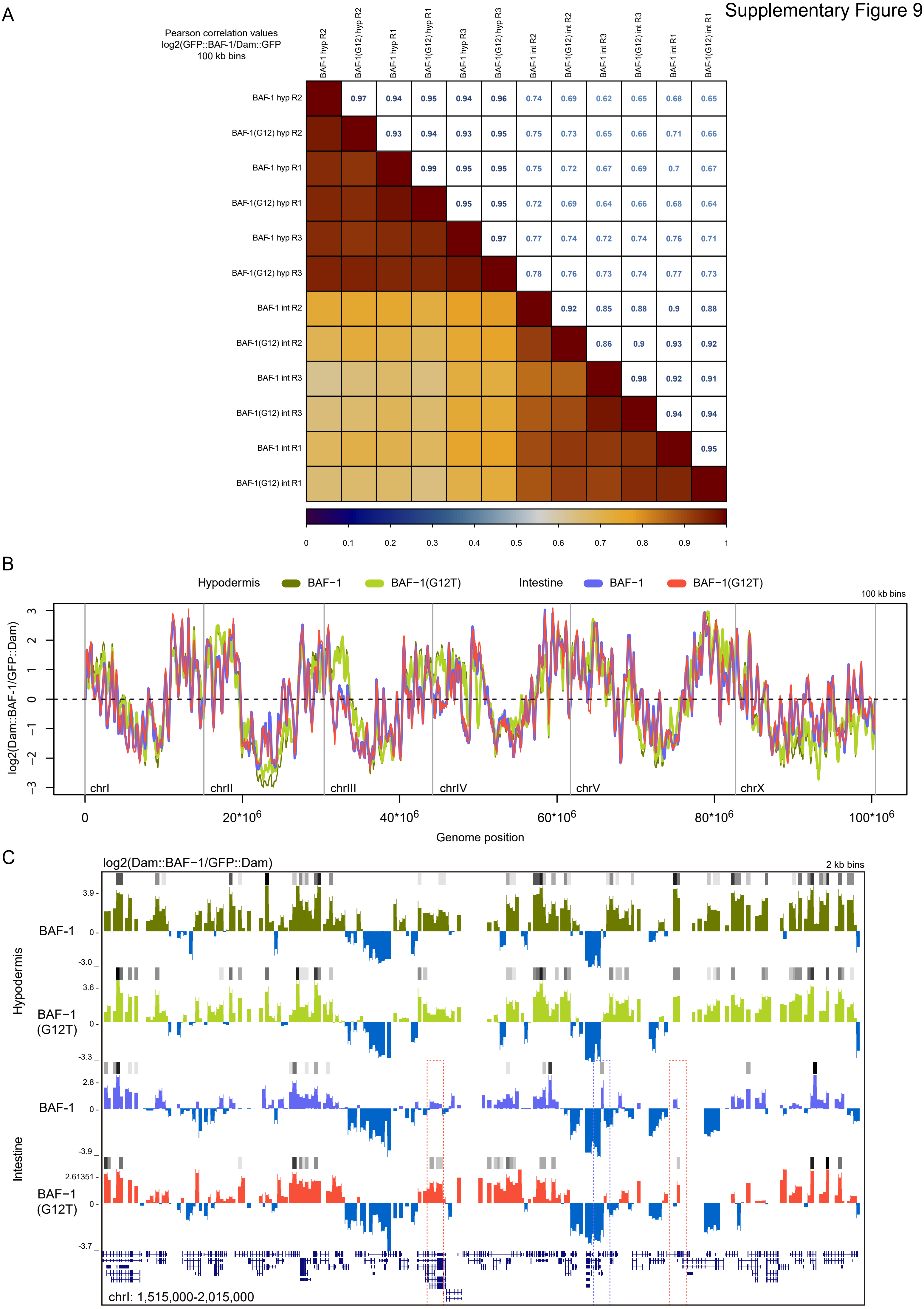

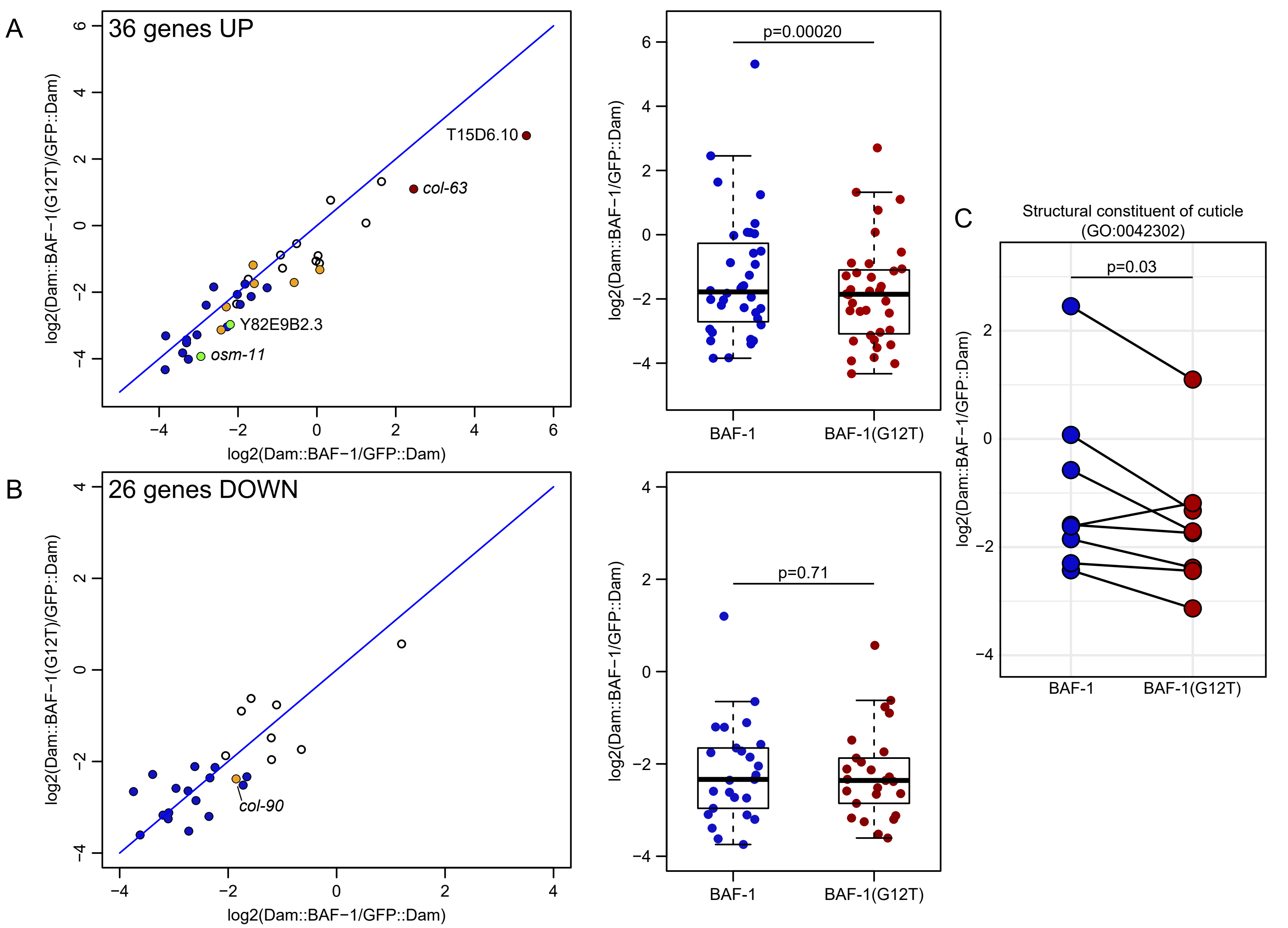
