## Supplementary Table S1 for "A progeria-associated BAF-1 mutation modulates gene expression and accelerates aging in *C. elegans*"

**Supplementary Table S1A**

| GFP::LMN-1 | | | | | | |
|---|---|---|---|---|---|---|
| | Day 1 | | Day 6 | | Day 8 | |
| Class | WT | *baf-1(G12T)* | WT | *baf-1(G12T)* | WT | *baf-1(G12T)* |
| I | 35 | 21 | 9 | 9 | 1 | 8 |
| II | 64 | 84 | 45 | 32 | 47 | 45 |
| III | 11 | 18 | 127 | 141 | 64 | 114 |
| IV | 2 | 0 | 21 | 55 | 9 | 40 |
| Total | 112 | 123 | 202 | 237 | 121 | 207 |
| Fisher's Exact Test p-values comparing WT and baf-1(G12T) | | | | | | |
| Class I vs II+III+IV | 0.029 | | | | | |
| Class I+II vs III+IV | | | 0.020 | | 0.0093 | |
| Class I+II+III vs IV | | | 0.00038 | | 0.0036 | |

**Supplementary Table S1B**

| EMR-1::mCherry | | | | | | |
|---|---|---|---|---|---|---|
| | Day 1 | | Day 6 | | Day 8 | |
| Class | WT | *baf-1(G12T)* | WT | *baf-1(G12T)* | WT | *baf-1(G12T)* |
| I | 29 | 27 | 25 | 1 | 0 | 5 |
| II | 38 | 72 | 87 | 16 | 43 | 19 |
| III | 3 | 17 | 151 | 202 | 219 | 177 |
| IV | 0 | 1 | 4 | 10 | 9 | 21 |
| Total | 70 | 117 | 267 | 229 | 271 | 222 |
| Fisher's Exact Test p-values comparing WT and baf-1(G12T) | | | | | | |
| Class I vs II+III+IV | 0.013 | | | | | |
| Class I+II vs III+IV | | | < 2.20E-16 | | 0.11 | |
| Class I+II+III vs IV | | | 0.062 | | 2.53E-06 | |

**Supplementary Table S1C**

| Combined | | | | | | |
|---|---|---|---|---|---|---|
| | Day 1 | | Day 6 | | Day 8 | |
| Class | WT | *baf-1(G12T)* | WT | *baf-1(G12T)* | WT | *baf-1(G12T)* |
| I | 64 | 48 | 34 | 10 | 1 | 13 |
| II | 102 | 156 | 132 | 48 | 90 | 64 |
| III | 14 | 35 | 278 | 343 | 283 | 291 |
| IV | 2 | 1 | 25 | 65 | 18 | 61 |
| Total | 182 | 240 | 469 | 466 | 392 | 429 |
| Fisher's Exact Test p-values comparing WT and baf-1(G12T) | | | | | | |
| Class I vs II+III+IV | 0.00055 | | | | | |
| Class I+II vs III+IV | | | < 2.20E-16 | | 0.069 | |
| Class I+II+III vs IV | | | 7.21E-06 | | 2.53E-06 | |

Classification of hypodermal nuclei expressing GFP::LMN-1 (A) or EMR-1::mCherry (B) according to their morphology (see Material and Methods). Combined data from (A) and (B) are represented in (C).
