## Supplementary Table S2 for "A progeria-associated BAF-1 mutation modulates gene expression and accelerates aging in *C. elegans*"

| | | Arm | Center | Ratio arm/center | p-value[a] |
|---|---|---|---|---|---|
| Hypodermis | All autosome bins | 5133 | 3125 | 1.6 | |
| | Enriched WT[b] | 358 | 52 | 6.9 | < 2.20E-16 |
| | Enriched G12T[c] | 228 | 250 | 0.9 | 5.03E-10 |
| Intestine | Enriched WT[b] | 320 | 266 | 1.2 | 0.00036 |
| | Enriched G12T[c] | 351 | 246 | 1.4 | 0.106 |

[a] Fisher's Exact Test p-values

[b] Bin enriched for WT (FDR <0.05; fold change >2) and that fulfil log2(Dam::BAF-1/GFP::Dam) - log2(Dam::BAF-1(G12T)/GFP::Dam) >0.58

[c] Bin enriched for G12T (FDR <0.05; fold change >2) and that fulfil log2(Dam::BAF-1(G12T)/GFP::Dam) - log2(Dam::BAF-1/GFP::Dam) >0.58
