## Supplementary Table S5 for "A progeria-associated BAF-1 mutation modulates gene expression and accelerates aging in *C. elegans*"

| Strain | Description | Genotype | Method | Reference |
|---|---|---|---|---|
| BN148 | Ubiquitous expression of EMR-1::mCh from integrated single-copy transgene; rescues *emr-1(gk119)* mutation | *emr-1(gk119) I; bqSi143[emr-1p::emr-1::mCh] II* | MosSCI | (Morales-Martinez et al. 2015) |
| BN189 | Ubiquitous expression of mCh::HIS-58 from integrated single-copy transgene | *bqSi189[lmn-1p::mCh::his-58)] II* | MosSCI | (Gomez-Saldivar et al. 2016) |
| BN448 | For tissue-specific control DamID after crossing to FLP driver | *bqSi447[hsp16.41p::FRT::mCh::his-58::FRT::gfp::dam] II* | MosSCI | (Cabianca et al. 2019)) |
| BN536 | For tissue-specific BAF-1 DamID after crossing to FLP driver | *bqSi536[hsp16.41p>mCh::his-58>dam::baf-1] II* | MosSCI | This study |
| BN548 | Expression of FLP in hypodermis | *bqSi548[dpy-7p::FLP] IV* | MosSCI | (Munoz-Jimenez et al. 2017) |
| BN561 | Control DamID in hypodermis | *bqSi447[hsp16.41p::FRT::mCh::his-58::FRT::gfp::dam] II; bqSi548[dpy-7p::FLP] IV* | MosSCI | (Fragoso-Luna et al. 2023) |
| BN580 | GFP knock-in into 5'-end of *baf-1* CDS; Frt sites in 1st and 2nd intron of GFP; designed for tissue-specific gene knockout | *baf-1(bq12[g>f>p::baf-1]) III* | CRISPR | (Munoz-Jimenez et al. 2017) |
| BN581 | mCh knock-in into 5'-end of *baf-1* CDS | *baf-1(bq13[mCh::baf-1]) III* | CRISPR | (Fragoso-Luna et al. 2023) |
| BN777 | For tissue-specific BAF-1(G12T) DamID after crossing to FLP driver and *baf-1(G12T)* | *bqSi777[hsp16.41p>mCh::his-58>dam::baf-1(G12T)] II* | MosSCI | This study |
| BN808 | *baf-1(G12T)* mutant | *baf-1(bq19[G12T]) III* | PMX375 crossed to N2 6 times | This study |
| BN868 | *baf-1(G12T)* mutant; uncoordinated | *baf-1(bq19[G12T]) unc-119(ed3) III* | BN808 crossed with HT1593 | This study |
| BN869 | Ubiquitous expression of mCh::HIS-58 and GFP::LMN-1 | *lmn-1(yc32[gfp::lmn-1]) I; bqSi189[lmn-1p::mCh::his-58] II* | BN189 crossed with UD484 | This study |
| BN870 | *baf-1(G12T)* mutant with ubiquitous expression of mCh::HIS-58 and heterozygous expression of GFP::LMN-1 | *lmn-1(yc32[gfp::lmn-1])/+ I; bqSi189[lmn-1p::mCh::his-58)] II; baf-1(bq19[G12T]) III* | BN808 crossed with BN189 and UD484 | This study |
| BN922 | *baf-1(G12T)* mutant with ubiquitous expression of EMR-1::mCh | *emr-1(gk119) I; bqSi143[emr-1p::emr-1::mCh] II; baf-1(bq19[G12T]) III* | BN148 crossed with BN808 | This study |
| BN998 | Expression of FLP in intestine | *bqSi997[nhx-2p::FLP::SL2::mNG] IV* | MosSCI | (Fragoso-Luna et al. 2023) |
| BN1007 | GFP knock-in into 5'-end of *baf-1(G12T)* CDS; Frt sites in 1st and 2nd intron of GFP; balanced w/ mT1 | *+/mT1 [umnIs34] II; baf-1(bq24[G>F>P::baf-1[G12T]])/mT1 [dpy-10(e128)] III* | CRISPR | This study |
| BN1024 | mCh knock-in into 3'-end of *emr-1* CDS; also Frt site in 5'-end of *emr-1* CDS | *emr-1(bq34[emr-1::mCh]) I* | CRISPR | This study |
| BN1037 | mCh knock-in into 3'-end of *emr-1* CDS; GFP knock-in into 5'-end of *lmn-1* CDS | *emr-1(bq34[emr-1::mCh]) lmn-1(yc32[gfp::lmn-1]) I* | BN1024 crossed with UD484 | This study |
| BN1047 | BAF-1 DamID in hypodermis | *bqSi536[hsp16.41p>mCh::his-58>dam::baf-1] II; bqSi548[dpy-7p::FLP] IV* | BN536 crossed with BN548 | This study |
| BN1048 | Control DamID in hypodermis of *baf-1(G12T)* | *bqSi447[hsp16.41p::FRT::mCh::his-58::FRT::gfp::dam] II; baf-1(bq19[G12T]) III; bqSi548[dpy-7p::FLP] IV* | BN561 crossed with BN808 | This study |
| BN1050 | BAF-1(G12T) DamID in hypodermis | *bqSi777[hsp16.41p>mCh::his-58>dam::baf-1(G12T)] II; baf-1(bq19[G12T]) III; bqSi548[dpy-7p::FLP] IV* | BN548 crossed with BN777 and BN808 | This study |
| BN1051 | Control DamID in intestine | *bqSi447[hsp16.41p>mCh::his-58>gfp::dam] II; bqSi997[nhx-2p::FLP::SL2::mNG] IV* | BN448 crossed with BN998 | This study |
| BN1052 | BAF-1 DamID in intestine | *bqSi536[hsp16.41p>mCh::his-58>dam::baf-1] II; bqSi997[nhx-2p::FLP::SL2::mNG] IV* | BN536 crossed with BN998 | This study |
| BN1053 | Control DamID in intestine of *baf-1(G12T)* | *bqSi447[hsp16.41p>mCh::his-58>gfp::dam] II; baf-1(bq19[G12T]) III; bqSi997[nhx-2p::FLP::SL2::mNG] IV* | BN808 crossed with BN1051 | This study |
| BN1054 | BAF-1(G12T) DamID in intestine | *bqSi777[hsp16.41p>mCh::his-58>dam::baf-1(G12T)] II; baf-1(bq19[G12T]) III; bqSi997[nhx-2p::FLP::SL2::mNG] IV* | BN777 crossed with BN808 and BN998 | This study |
| BN1150 | mCh knock-in into 3'-end of emr-1 CDS and GFP knock-in into 5'-end of lmn-1 CDS | *emr-1(bq34[emr-1::mCh]) lmn-1(yc32[gfp::lmn-1]) I* | BN1024 crossed with UD484 and then with N2 4 times | This study |
| BN1188 | GFP knock-in into 5'-end of *baf-1(G12T)* CDS; ubiquitous expression of mCh::HIS-58; balanced w/ mT1 | *bqSi189[lmn-1p::mCherry::his-58]/mT1 [umnIs34] II; baf-1(bq24[G>F>P::baf-1[G12T]])/mT1 [dpy-10(e128)] III* | BN189 crossed with BN1007 | This study |
| BN1337 | Balanced *baf-1(G12T)* mutant with ubiquitous expression of mCh::HIS-58 and heterozygous expression of GFP::LMN-1 | *lmn-1(yc32[gfp::lmn-1]) I/hT2 (I;III); bqSi189[lmn-1p::mCherry::his-58] II; baf-1(bq19[G12T]) III/hT2 (I;III)* | BN870 crossed with hT2 strain | This study |
| BN1349 | mCh knock-in into 5'-end of *baf-1* CDS; temperature-sensitive; at 25C animals lack germ line | *glp-4(bn2) I; baf-1(bq13[mCh::baf-1]) III* | BN581 crossed with SS104 | This study |
| BN1350 | *baf-1(G12T)* mutant; temperature-sensitive; at 25C animals lack germ line | *glp-4(bn2) I; baf-1(bq19[G12T]) III* | BN808 crossed with BN1349 | This study |
| BN1375 | Temperature-sensitive; at 25C animals lack germline | *glp-4(bn2) I* | BN808 crossed with BN1349 | This study |
| BN1412 | Tissue-specific RPB-6 DamID | *bqSi1411[hsp16.41p::FRT::mCh::his-58::FRT::dam::rpb-6] II; bqSi577[myo-2p::GFP)] IV* | MosSCI | (Fragoso-Luna et al. 2023) |
| BN1414 | RPB-6 DamID in hypodermis | *bqSi1411[hsp16.41p::FRT::mCh::his-58::FRT::dam::rpb-6] II; bqSi548[dpy-7p::FLP] IV* | MosSCI | (Fragoso-Luna et al. 2023) |

| | | | | |
|---|---|---|---|---|
| BN1415 | RPB-6 DamID in intestine | *bqSi1411[hsp16.41p::FRT::mCh::his-58::FRT::dam::rpb-6] II; bqSi997[nhx-2p::FLP::SL2::mNG] IV* | BN998 crossed with BN1412 | This study |
| BN1416 | RPB-6 DamID in hypodermis in *baf-1(G12T)* | *bqSi1411[hsp16.41p::FRT::mCh::his-58::FRT::dam::rpb-6] II; baf-1(bq19[G12T]) III; bqSi548[dpy-7p::FLP] IV* | BN808 crossed with BN1414 | This study |
| BN1417 | RPB-6 DamID in intestine in *baf-1(G12T)* | *bqSi1411[hsp16.41p::FRT::mCh::his-58::FRT::dam::rpb-6] II; baf-1(bq19[G12T]) III; bqSi997[nhx-2p::FLP::SL2::mNG)] IV* | BN808 crossed with BN1415 | This study |
| CB4108 | Feminized hermaphrodites unable to producing sperm | *fog-2(q71) V* | | (Katju et al. 2008) |
| EG4322 | MosSCI targeting strain | *ttTi5605 II; unc-119(ed9) III* | | (Katju et al. 2008) |
| HT1593 | Uncoordinated | *unc-119(ed3) III* | | Dickinson et al. 2013) |
| N2 | *C. elegans* var Bristol | | | CGC |
| PMX375 | *baf-1(G12T)* mutant | *baf-1(bq19[G12T]) III* | CRISPR | This study |
| SS104 | Temperature-sensitive; at 25C animals lack germline | *glp-4(bn2) I* | | (Beanan & Strome, 1992) |
| UD484 | GFP knock-in into 5'-end of *lmn-1* CDS | *lmn-1(yc32[gfp::lmn-1]) I* | CRISPR | (Bone et al. 2016) |
| UV117 | GFP knock-in into 5'-end of *lmn-1* CDS | *lmn-1(jf98[lmn-1::GFP]) I* | CRISPR | (Link et al. 2018) |

Strains used in this study
