## Supplementary Table S6 for "A progeria-associated BAF-1 mutation modulates gene expression and accelerates aging in *C. elegans*"

| Primer ID | Description | Sequence |
|---|---|---|
| B633 | generation of pBN180 | 5'-AACGTCGTGACTGGGAAAAC |
| B634 | generation of pBN180 | 5'-GGTGCCAACTTTTCTATACAAAG |
| B635 | universal sgRNA primer | 5'-CAAGACATCTCGCAATAGGAG |
| B724 | dpy-10; sgRNA sequence in capitals | 5'-CTACCATAGGCACCACGAGgttttagagctagaaatagcaagt |
| B725 | dpy-10; repair template | 5'-<br>CACTTGAACTTCAATACGGCAAGATGAGAATGACTGGAAACCGTACCGCATG<br>CGGTGCCTATGGTAGCGGAGCTTCACATGGCTTCAGACCAACAGCCTAT |
| B780 | generation of pBN215 | 5'-aaacagcatagcaagtttAAATAAGGCTAGTCCGTTATC |
| B781 | generation of pBN215 | 5'-ccagcatagctcttaaacCAAGACATCTCGCAATAG |
| B825 | emr-1 3'-end; sgRNA sequence in capitals | 5'-CGTGGCCGAGACGAATCCGGgtttaagagctatgctggaaac |
| B838 | emr-1 5'-end; sgRNA sequence in capitals | 5'-AGACGGAAGCATTAACAGTGgtttaagagctatgctggaaac |
| B840 | emr-1::mCh gBlock w/ Frt sites | 5'-<br>TGGGCCAATTGTGGCGACGACCCGCAAGCTCTACGAGAAGAAGCTTATCAA<br>GTTATCAGAtGGAAGCATTAACAgtgagCatttgaatttaaattatttattaatttctGAAGTTCCT<br>ATTCTCTAGAAAGTATAGGAACTTCaaataacatttttaaagttttcgaacctctaatttccagaTC<br>AATCAAATCTCAACGACTCTCAATTCAACGAGGATTCATTGATCATCAGCTCG<br>TCACCGAAGAAATCACCGCCACAACGAGTTTTCCAGAACGTGTCAGCTGCAA<br>CAGCGGCAGCTACTACCTCTCCCGAATCGGACAGCGACGATTGCGAGGAGT<br>CGATGCGATACTTGACGGAAGAGGAAATGGCCGCCGATCGGGCATCGGCTC<br>GTCAAGCTCAGAGCAACAAAGGAGGATTCTTGGGAAGCACGgtgagtttttcgcccttttt<br>tctggcaataaaaacttattttctattttccagATCACATTCACAATTCTCTTCGTCTTCATCGCC<br>GTCTTCGCCTACTTCTTGATCGAGAACGCCGAGCAGTTGAAGCTCGTGGCCG<br>AGACGAATCCaGAaGATACTATTtgtacaGTCTCAAAGGGTGAAGAAGATAACAT<br>GGCAATTATTAAAGAGTTTATGCGTTTCAAGGTGCATATGGAGGGATCTGTCA<br>ATGGGCATGAGTTTGAAATTGAAGGTGAAGGAGAAGGCCGACCATATGAGG<br>GAACACAAACCGCAAAACTAAAGgtaagtttaaacatatatatGAAGTTCCTATTCTCTA<br>GAAAGTATAGGAACTTCactaactaaccctgattatttaaattttcagGTAACTAAAGGCGGAC<br>CATTACCATTCGCCTGGGACATCCTCTCTCCACAGTTCATGTATGGAAGTAAA<br>GCTTATGTTAAACATCCGGCAGATATACCAGATTATTTGAAACTTTCATTCCC<br>GGAGGGTTTTAAGTGGGAACGCGTAATGA |
| B914 | AdRt | 5'-CTAATACGACTCACTATAGGGCAGCGTGGTCGCGGCCGAGGA |
| B919 | AdRb | 5'-TCCTCGGCCGCG |
| B920 | AdR | 5'-NNNNGTGGTCGCGGCCGAGGATC |
| B925 | baf-1 5'-end | 5'-caagatctATGTCGACTTCTGTTAAGCATCG |
| B926 | baf-1 3'-end | 5'-tagctagcTTACATGAACTGATCTGCCC |
| B1436 | AdR4N | 5'-NNNNGTCCTCGCGGCCGAGGATC |
| B1437 | AdR5N | 5'-NNNNNGTCCTCGCGGCCGAGGATC |
| B1438 | AdR6N | 5'-NNNNNNGTCCTCGCGGCCGAGGATC |
| B1103 | pmp-3 qRT-PCR | 5'-TGGCCGGATGATGGTGTCGC |
| B1104 | pmp-3 qRT-PCR | ACGAACAATGCCAAAGGCCAGC |
| B1105 | tba-1 qRT-PCR | TCAACACTGCCATCGCCGCC |
| B1106 | tba-1 qRT-PCR | TCCAAGCGAGACCAGGCTTCAG |
| B1107 | Y45F10D.4 qRT-PCR | CGAGAACCCGCGAAATGTCGGA |
| B1108 | Y45F10D.4 qRT-PCR | CGGTTGCCAGGGAAGATGAGGC |
| B1603 | baf-1 qRT-PCR | ATGCAGGCTTCGATAAAGCCTAC |
| B1604 | baf-1 qRT-PCR | CAGCCGTCTCTTTCAGCCAT |
| B1605 | emr-1 qRT-PCR | GCATTAACAATCAATCAAATCTC |
| B1606 | emr-1 qRT-PCR | AAAACTCGTTGTGGCGGTGA |
| B1607 | lmn-1 qRT-PCR | TGGTGGTGGAGAATGATGATCTC |
| B1608 | lmn-1 qRT-PCR | CGGCTGTTCGGAGAAGAGTT |

Primers used in this study
