## Supplementary Table S7 for "A progeria-associated BAF-1 mutation modulates gene expression and accelerates aging in *C. elegans*"

| Sample | Dam fusion | *baf-1* | Tissue | Experiment | Raw.reads | Cut.reads | Mapped.reads | GATC.reads | Percentage |
|---|---|---|---|---|---|---|---|---|---|
| BN1047R1 | Dam::BAF-1 | WT | Hyp | BAF-1 DamID | 18740263 | 3005144 | 1485068 | 1481972 | 7.9 |
| BN1047R2 | Dam::BAF-1 | WT | Hyp | BAF-1 DamID | 23215175 | 9272778 | 2350374 | 2322595 | 10.0 |
| BN1047R3 | Dam::BAF-1 | WT | Hyp | BAF-1 DamID | 27642934 | 9395426 | 2982162 | 2933349 | 10.6 |
| BN561R1 | GFP::Dam | WT | Hyp | BAF-1 DamID | 20511930 | 3382859 | 1982157 | 1979881 | 9.7 |
| BN561R2 | GFP::Dam | WT | Hyp | BAF-1 DamID | 11999338 | 3534312 | 1847526 | 1838635 | 15.3 |
| BN561R3 | GFP::Dam | WT | Hyp | BAF-1 DamID | 6835704 | 2211321 | 1473010 | 1466550 | 21.5 |
| BN1052R1 | Dam::BAF-1 | WT | Int | BAF-1 DamID | 39808838 | 7620330 | 896733 | 890652 | 2.2 |
| BN1052R2 | Dam::BAF-1 | WT | Int | BAF-1 DamID | 58252001 | 9750214 | 536843 | 515492 | 0.9 |
| BN1052R3 | Dam::BAF-1 | WT | Int | BAF-1 DamID | 40395718 | 6985997 | 4653416 | 4609737 | 11.4 |
| BN1051R1 | GFP::Dam | WT | Int | BAF-1 DamID | 11620807 | 1957742 | 1466782 | 1465633 | 12.6 |
| BN1051R2 | GFP::Dam | WT | Int | BAF-1 DamID | 66542014 | 12815500 | 8997154 | 8973703 | 13.5 |
| BN1051R3 | GFP::Dam | WT | Int | BAF-1 DamID | 39093117 | 5577439 | 4257133 | 4242468 | 10.9 |
| BN1050R1 | Dam::BAF-1(G12T) | *baf-1(G12T)* | Hyp | BAF-1 DamID | 13500562 | 3095446 | 1517815 | 1513434 | 11.2 |
| BN1050R2 | Dam::BAF-1(G12T) | *baf-1(G12T)* | Hyp | BAF-1 DamID | 28347252 | 8574994 | 1713384 | 1629751 | 5.7 |
| BN1050R3 | Dam::BAF-1(G12T) | *baf-1(G12T)* | Hyp | BAF-1 DamID | 10603157 | 3537523 | 1730454 | 1719304 | 16.2 |
| BN1048R1 | GFP::Dam | *baf-1(G12T)* | Hyp | BAF-1 DamID | 14039486 | 2459948 | 1836312 | 1833763 | 13.1 |
| BN1048R2 | GFP::Dam | *baf-1(G12T)* | Hyp | BAF-1 DamID | 19416940 | 4604504 | 2860379 | 2847206 | 14.7 |
| BN1048R3 | GFP::Dam | *baf-1(G12T)* | Hyp | BAF-1 DamID | 13719184 | 4591705 | 3150279 | 3143647 | 22.9 |
| BN1054R1 | Dam::BAF-1(G12T) | *baf-1(G12T)* | Int | BAF-1 DamID | 45560629 | 6473550 | 2889857 | 2866010 | 6.3 |
| BN1054R2 | Dam::BAF-1(G12T) | *baf-1(G12T)* | Int | BAF-1 DamID | 55998615 | 7228206 | 3259482 | 2998322 | 5.4 |
| BN1054R3 | Dam::BAF-1(G12T) | *baf-1(G12T)* | Int | BAF-1 DamID | 10156315 | 3489919 | 2179779 | 2159861 | 21.3 |
| BN1053R1 | GFP::Dam | *baf-1(G12T)* | Int | BAF-1 DamID | 10397627 | 2119217 | 1533011 | 1531288 | 14.7 |
| BN1053R2 | GFP::Dam | *baf-1(G12T)* | Int | BAF-1 DamID | 6735160 | 1825474 | 1154150 | 1150780 | 17.1 |
| BN1053R3 | GFP::Dam | *baf-1(G12T)* | Int | BAF-1 DamID | 45604517 | 5500716 | 3849699 | 3842500 | 8.4 |
| BN1414R1 | Dam::RPB-6 | WT | Hyp | RAPID | 55151578 | 49896285 | 7327585 | 7259801 | 13.2 |
| BN1414R2 | Dam::RPB-6 | WT | Hyp | RAPID | 29472050 | 27283523 | 11135772 | 11125581 | 37.7 |
| BN561R1 | GFP::Dam | WT | Hyp | RAPID | 25851173 | 23996691 | 15318783 | 15312243 | 59.2 |
| BN561R2 | GFP::Dam | WT | Hyp | RAPID | 34142422 | 31912638 | 19855436 | 19849763 | 58.1 |
| BN1415R1 | Dam::RPB-6 | WT | Int | RAPID | 38725533 | 36232658 | 10573724 | 10567462 | 27.3 |
| BN1415R2 | Dam::RPB-6 | WT | Int | RAPID | 51270952 | 46191739 | 15459449 | 15420825 | 30.1 |
| BN1051R1 | GFP::Dam | WT | Int | RAPID | 27495129 | 25830819 | 15911235 | 15907256 | 57.9 |
| BN1051R2 | GFP::Dam | WT | Int | RAPID | 24825994 | 23123070 | 13559944 | 13556832 | 54.6 |
| BN1416R1 | Dam::RPB-6 | *baf-1(G12T)* | Hyp | RAPID | 76541282 | 68788560 | 3552440 | 3430318 | 4.5 |
| BN1416R2 | Dam::RPB-6 | *baf-1(G12T)* | Hyp | RAPID | 107439981 | 100096170 | 27493236 | 27451137 | 25.6 |
| BN1048R1 | GFP::Dam | *baf-1(G12T)* | Hyp | RAPID | 22861824 | 21200127 | 12544566 | 12538120 | 54.8 |
| BN1048R2 | GFP::Dam | *baf-1(G12T)* | Hyp | RAPID | 28267352 | 26364680 | 16859072 | 16853186 | 59.6 |
| BN1417R1 | Dam::RPB-6 | *baf-1(G12T)* | Int | RAPID | 32389256 | 30146272 | 10307381 | 10294746 | 31.8 |
| BN1417R2 | Dam::RPB-6 | *baf-1(G12T)* | Int | RAPID | 33315508 | 30389405 | 10086027 | 10016998 | 30.1 |
| BN1053R1 | GFP::Dam | *baf-1(G12T)* | Int | RAPID | 30530830 | 28777590 | 17602729 | 17598709 | 57.6 |
| BN1053R2 | GFP::Dam | *baf-1(G12T)* | Int | RAPID | 39309242 | 36788497 | 21988543 | 21983661 | 55.9 |

Individual DamID samples obtained in this study
